# The Hippo pathway oncoprotein YAP promotes melanoma cell invasion and spontaneous metastasis

**DOI:** 10.1101/835454

**Authors:** Xiaomeng Zhang, Lie Yang, Pacman Szeto, Youfang Zhang, Kaushalya Amarasinghe, Jason Li, Catriona McLean, Mark Shackleton, Kieran F. Harvey

## Abstract

Melanoma is a deadly form of skin cancer that accounts for a disproportionally large proportion of cancer-related deaths in younger people. Compared to most other skin cancers, a feature of melanoma is its high metastatic capacity, although molecular mechanisms that confer this are not well understood. The Hippo pathway is a key regulator of organ growth and cell fate that is deregulated in many cancers. To analyse the Hippo pathway in cutaneous melanoma, we generated a transcriptional signature of pathway activity in melanoma cells. Hippo-mediated transcriptional activity varied in melanoma cell lines but failed to cluster with known genetic drivers of melanomagenesis such as *BRAF* and *NRAS* mutation status. Instead, it correlated strongly with published gene expression profiles linked to melanoma cell invasiveness. Consistent with this, the central Hippo oncogene, YAP, was both necessary and sufficient for melanoma cell invasion in vitro. In in vivo murine studies, YAP promoted spontaneous melanoma metastasis, whilst the growth of YAP-expressing primary tumours was impeded. Finally, we identified the YAP target genes *AXL*, *THBS1* and *CYR61* as key mediators of YAP-induced melanoma cell invasion. These data suggest that the Hippo pathway is a critical regulator of melanoma metastasis.

## INTRODUCTION

Cutaneous melanoma is a deadly disease that has a disproportionately high socioeconomic impact because of its poor prognosis and propensity to present earlier in life than most other cancers. The majority of cutaneous melanomas are driven by hyperactivation of the MAPK pathway, with common mutations in the *BRAF*, *NRAS* and *NF1* genes (1, 2). The past decade has witnessed substantial improvements in melanoma treatment with the development of both MAPK pathway inhibitors and immunotherapies. MAPK pathway inhibitors have shown substantial activity in patients harbouring activating *BRAF* mutations, but resistance to these therapies is rapid in most cases (3–5). Immunotherapies, most notably those targeting PD-1 or its ligand, have shown dramatic sustained responses in many but not all patients (6, 7). As such, better therapeutic options for melanoma are required.

A feature of melanoma cells is phenotypic plasticity. Cell culture experiments revealed that melanoma cells can be broadly classified into at least two groups based on their gene expression profile, expressing genes associated with either invasive or proliferative behaviour that is observed in vitro (8–10). Invasive and proliferative melanoma cell states were subsequently shown to exist in vivo in human tumours (11). Based on bulk RNA-sequencing of melanomas, tumours were initially thought to possess cells expressing either the proliferative or the invasive signature. However, single cell RNA-sequencing studies subsequently showed that melanomas can comprise cells in either proliferative or invasive states, in differing ratios (12). Archetypal markers of these different states include the MITF transcription factor (proliferative state) and the AXL receptor tyrosine kinase (invasive state). Interestingly, numerous studies have reported that melanoma cells can transition between the invasive and proliferative states, rather than being locked into one state. Such changes are not driven by genetic mutations but are mediated by changes in the cellular transcriptome, downstream of signalling events. For example, treatment of melanoma cells with BRAF inhibitors (BRAFi) drives them towards a more drug-resistant state with low MITF and high AXL expression, reminiscent of the invasive melanoma state (13, 14). As well as MITF, the proliferative melanoma subtype is associated with expression of the SOX10 and PAX3 transcription factors. In contrast, the TEAD1-TEAD4 and AP-1 family transcription factors promote invasive melanoma subtypes (11). In independent studies, c-Jun (which is an AP-1 family transcription factor) was also shown to be essential for BRAFi resistance in melanoma (15, 16).

The TEAD1-TEAD4 transcription factors are key downstream mediators of the Hippo pathway and cooperate with the YAP and TAZ transcription co-activator proteins to regulate transcription (17–19). Together, YAP, TAZ and TEAD1-4 (and their respective *Drosophila* orthologues Yorkie and Scalloped) control organ growth and cell fate downstream of the Hippo pathway (17–19). Hippo limits activity of YAP, TAZ and Yorkie by controlling the rate at which they transit between the nucleus and cytoplasm (20–22). The Hippo pathway also regulates many cellular behaviours that underpin cancer such as cell proliferation, cell survival, metastasis and cell fate control (23). In addition, this pathway has been identified as a mediator of drug resistance, both chemotherapies and targeted therapies like MAPK inhibitors, in cutaneous melanoma and other MAPK-driven tumours (24–27).

Hippo pathway deregulation is common in many solid cancers like lung, breast and liver, whilst pathway mutations are infrequent (23). In certain cancers, such as mesothelioma and meningioma, mutation of Hippo pathway genes occurs in approximately 50% of cases and is considered a driving event (28–30). Additionally, this pathway is thought to be central to uveal melanomagenesis as YAP is hyperactive in uveal melanoma cells and mediates the oncogenic effect of *GNAQ* and *GNA11* mutations, which occur in the vast majority of these cancers (31, 32). By contrast, the role of Hippo/YAP signalling in cutaneous melanoma is less clear. Here, by generating a transcriptional signature of YAP in melanoma cells we find that YAP activity is elevated almost exclusively in the invasive class of melanoma cell lines. YAP is required for melanoma cell invasion and YAP hyperactivity can switch the melanoma cell phenotype from proliferative to invasive by driving expression of *AXL*, *CYR61* and *CRIM1*. Constitutive YAP hyperactivity promotes spontaneous melanoma metastasis in murine xenografts, but compromises primary tumour growth, possibly because it impedes melanoma cell plasticity.

## RESULTS

### YAP activity is elevated in invasive melanoma cell lines

Previously, we reported a variable requirement of YAP for melanoma cell viability across a panel of ten cell lines and three patient-derived xenografts, i.e. some cell lines and xenografts were highly dependent on YAP for survival whereas others were not (33). We observed no obvious correlation between sensitivity to YAP depletion and the major melanoma genotypes (*BRAF* mutant or *NRAS* mutant), and no correlation with YAP activity, as determined by YAP phosphorylation status at S127 (33). In addition, we observed no striking fluctuation in YAP expression or YAP nuclear/cytoplasmic ratio in different stages of melanomagenesis in humans, although YAP was overexpressed in most melanomas compared to normal melanocytes (33). As analysis of YAP target gene expression provides a more robust way to assess YAP activity, we sought to investigate YAP/Hippo’s role in cutaneous melanoma by identifying a YAP signature in melanoma cells. To do this we generated MeWo melanoma cells stably expressing Doxycycline (Dox)-inducible vectors (vector control or a hyperactive YAP allele, YAP-5SA). Cells were treated with Dox for 16 hours, and then total RNA was harvested and sequenced to identify differentially expressed genes immediately after YAP overexpression. 176 genes were significantly elevated in YAP-5SA-expressing cells (log2 fold change, p<0.05) and constituted a melanoma cell YAP signature, whilst 67 genes were downregulated (log2 fold change, p<0.05) (Table S1). Q-PCR was used to validate the expression of 13 genes (11 that were reported as being elevated in YAP-5SA-expressing cells and 2 that did not change). All 13 genes behaved similarly in both QPCR and RNA-seq studies, thus validating the RNA-seq experiments (Fig. S1). Among these genes were several validated YAP target genes (e.g. *CTGF*, *ITGB2* and *CYR61*). We also assessed expression of *AXL*, which has been identified as a YAP target gene (34). *AXL* was also elevated in YAP-5SA expressing MeWo cells, as determined by QPCR (Fig. S1).

We then assessed this melanoma YAP signature on gene expression profiling data from a panel of 55 melanoma cell lines (E-MTAB-1946) using unsupervised hierarchical clustering. This revealed two broad groups of cell lines: those with a YAP signature and those with low expression of YAP target genes (Fig. S2). We found no association between YAP activity and the mutation status of *BRAF* and *NRAS*, which are major melanoma driver genes (Fig. S2 and S3), consistent with our previous finding that YAP sensitivity in melanoma cells does not correlate with either *BRAF* or *NRAS* mutation status (33).

Previous studies identified two pervasive gene expression profiles in melanoma cell lines, which were termed invasive and proliferative (8) and subsequently shown to exist in patient tumours (11). Melanoma cells expressing an invasive signature displayed greater invasive properties in culture, whilst cells expressing a proliferative signature were less invasive in culture and tended to show higher proliferation rates and increased expression of genes associated with melanocytic cell differentiation, such as *MITF* (10). The invasive cell lines were also linked to MAPK inhibitor resistance and increased expression of the receptor tyrosine kinase *AXL* (13, 14).

We used the invasive signature defined by Hoek et al. (10) to perform unsupervised hierarchical clustering on gene expression data from the E-MTAB-1946 cell line panel and, consistent with previous studies on expression data from other melanoma cell line cohorts, identified two groups that either expressed the invasive or proliferative signatures (Fig. 1A). We then compared this with cell lines that we had identified as either YAP-high or YAP-low, based on levels of expression YAP target genes in our melanoma YAP signature. Strikingly, YAP-high cells almost exclusively belonged to the invasive group of melanoma cell lines (Fig. 1A).

**Figure 1.**
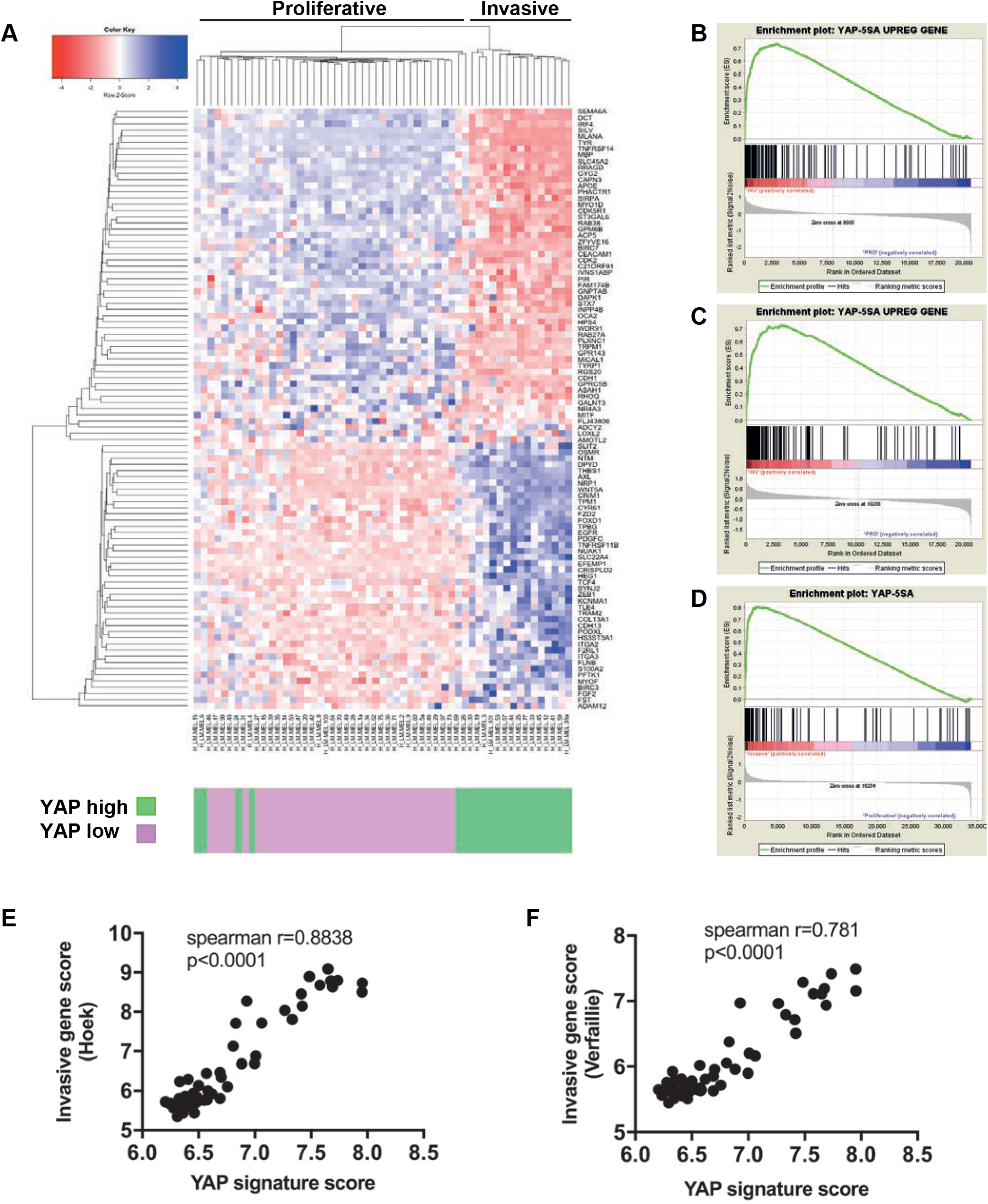
YAP activity is elevated in invasive melanoma cell lines. **A)** Heat map of mRNA expression of genes that constitute the Hoek melanoma invasion signature in a panel of melanoma cell lines. Unsupervised hierarchical clustering identified two groups (invasive or proliferative) with either YAP high or YAP low activity. YAP activity status is indicated below each cell line (YAP high cells are green and YAP low cells are pink). **B-D)** Gene set enrichment analysis plots of the YAP melanoma signature in three independent melanoma cell line gene expression databases: MTAB-1946 (**B**); GSEA4843 (**C**); and GSE1727(**D**). The YAP melanoma signature is enriched in cell lines invasive cell lines, as determined using the Hoek invasion signature. All enrichments were highly significant. **E-F)** Correlation analyses between the YAP melanoma signature and either the Hoek invasive signature (**E**) or the Verfaillie invasive signature (**F**). Both correlations were highly significant.

To investigate this further we performed gene set enrichment analysis on the E-MTAB-1946 cell line panel and found that the YAP signature was strongly enriched in invasive cell lines (Fig. 1B). We then performed gene set enrichment analysis on two additional melanoma cell line panels with associated microarray expression data (GSEA4843 and GSE1727) and found similarly strongly enrichment of the YAP signature in invasive cell lines (Fig. 1C and D). Finally, we performed pairwise analysis to compare the melanoma YAP signature to two signatures previously associated with invasive melanoma cell lines, the Hoek signature (10), and the Verfaillie signature (11). We observed a strong correlation between the melanoma YAP signature and both invasive signatures (Figure 1E and F). Collectively, these data show that YAP activity is substantially higher in melanoma cell lines that express an invasive gene expression profile.

### YAP can induce invasion in normally non-invasive melanoma cells

We and others previously linked YAP to migratory and invasive behaviour of cultured cells (35, 36), and so we investigated this in melanoma cell lines. Initially, we tested melanoma cell lines for their intrinsic ability to invade through a semi-porous membrane lined with matrigel to a chemoattractant and found that all cell lines fell into two groups: invasive and non-invasive (Fig. S4). We then expressed Doxycycline (Dox)-inducible YAP-5SA or control empty vector in two non-invasive cell melanoma lines and assessed their ability to invade in the above assays. MeWo or C013-M1 vector control cells treated with or without Dox, or untreated YAP-5SA cells, invaded poorly. In contrast, expression of YAP-5SA for 24 hours strongly stimulated the invasive capacity of both lines (Fig. 2A and B). This observation was independent of potential effects of YAP-5SA on cell number, as this was not significantly different between these conditions in the time course of this experiment (Fig. 2C and D). YAP expression levels in each experimental situation were determined using a YAP antibody (Fig. 2E). Similar results were obtained from a related cell migration assay (using a semi-porous membrane without matrigel), where YAP-5SA expression induced substantial migration of MeWo cells, compared to control cells (Fig S5). This demonstrates that expression of hyperactive YAP is sufficient to induce invasive behaviour in normally non-invasive melanoma cells.

**Figure 2.**
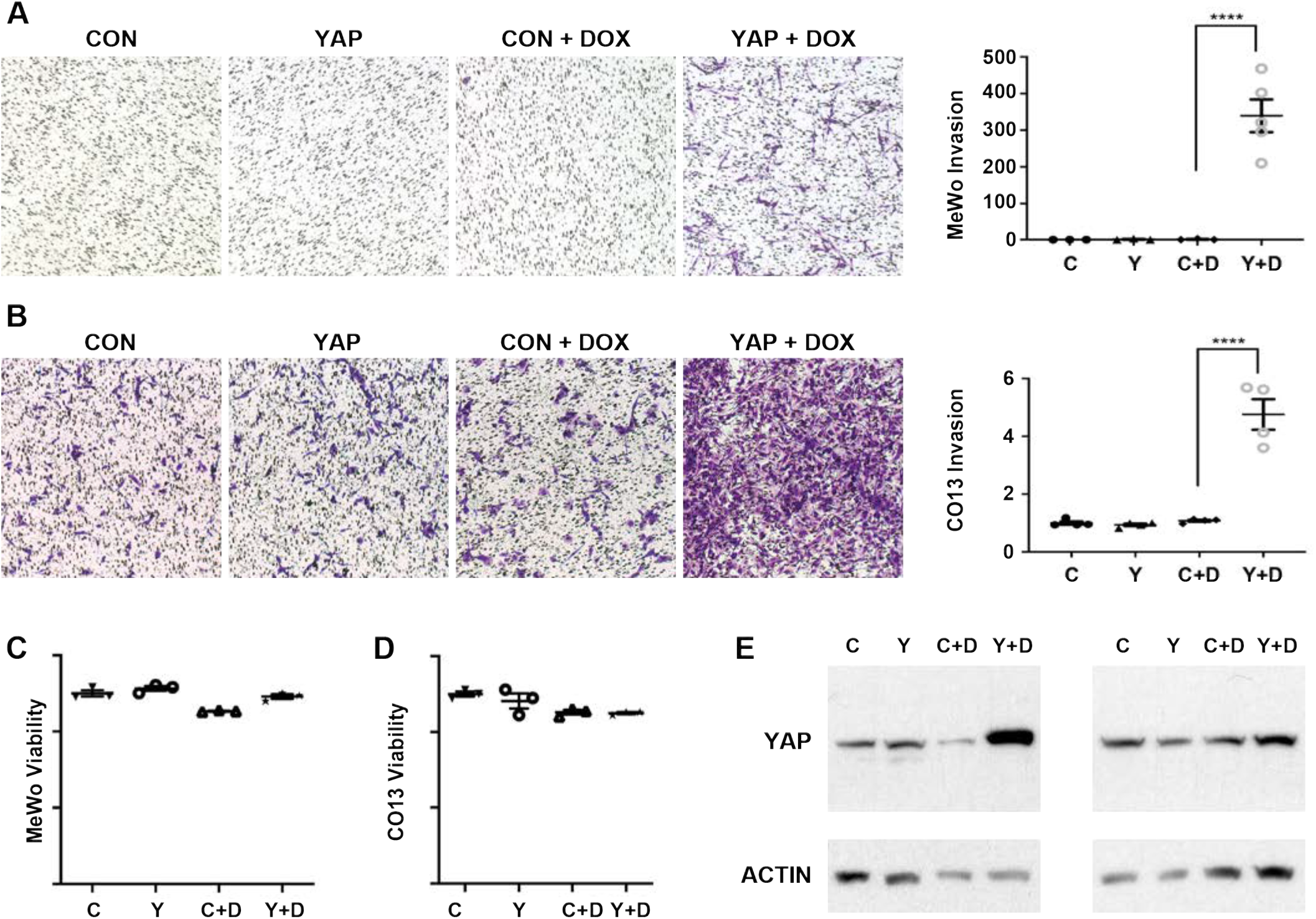
YAP can induce invasion in normally non-invasive melanoma cells. **A and B)** Representative images of melanoma cell lines following invasion assays and quantification of these assays (far right panel). MeWo cells (A) or C013-M1 cells (B) expressed YAP-5SA or control plasmids and were untreated or treated with Dox, as indicated. **C and D)** Quantification of viability assays for the different cell lines and treatments in (A) and (B). MeWo cells are plotted in (C) and C013-M1 cells in (D). **E)** Detection of YAP and actin by western blot for the different cell lines and treatments in (A) and (B). The left panel shows data from MeWo cells and the right panel is from C013-M1 cells. Data in (A-D) are represented as mean +/- SEM from 3 biological replicates. ****p<0.0001 (unpaired two-tailed t-tests or ANOVA plus Tukey’s multiple comparison tests).

### YAP is required for the invasive ability of melanoma cells

To determine whether YAP is required for the invasive ability of cells that express an invasive gene expression signature, we performed loss of function studies in C067-M1, A375 and HMCB cells, which we found to possess inherent invasive activity (Fig. S4). Each cell line was treated with YAP siRNA or control siRNA for 48 hours and subjected to invasion assays over 24 hours. YAP depletion significantly impaired the ability of these lines to invade (Fig. 3A-C). Again, these observations were independent of any effects of YAP depletion on cell number (Fig. 3A-C). Similarly, YAP depletion significantly inhibited the migratory ability of C067-M1 cells in cell migration assays (Fig. 3D). YAP depletion by siRNA levels in each cell line was confirmed using a YAP antibody (Fig. 3E). This shows that YAP is necessary for the invasive properties of melanoma cell lines that are inherently invasive in vitro.

**Figure 3.**
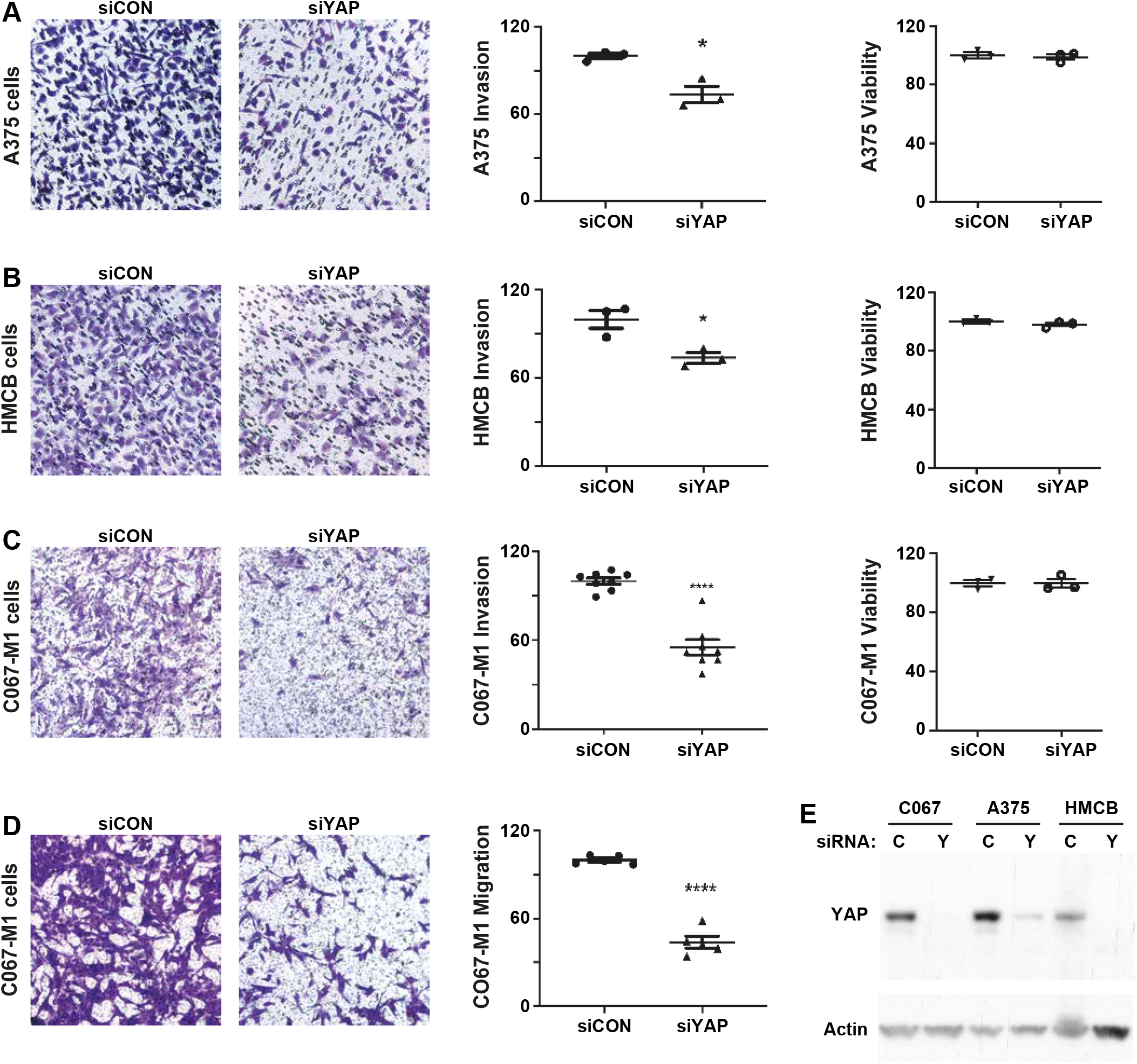
YAP is required for the invasive ability of melanoma cells. **A-C)** Representative images of the indicated melanoma cell lines following invasion assays. Cells were transfected with either control siRNA or YAP siRNA. Invasion and viability of these cells were quantified. Cell lines were A375 (A), HMCB (B) and C067-M1 (C). **D)** Representative images of C067-M1 cells transfected with either control siRNA or YAP siRNA and following migration assays. **E)** Detection of YAP and actin by western blot for the different cell lines and siRNA treatments in (A - C). Data in (A-D) are represented as mean +/- SEM from 3 biological replicates. *p<0.05, ****p<0.0001 (unpaired two-tailed t-tests or ANOVA plus Tukey’s multiple comparison tests).

### YAP induces spontaneous melanoma metastasis in vivo

Based on the above, we predicted that YAP might stimulate spontaneous melanoma cell metastasis from primary tumours in vivo. Previously, YAP has been studied in melanoma cell lines grown in mice following injection into the tail vein (37, 38). Although these studies suggested a role for YAP in metastasis, a major limitation of such experiments is that they do not assess the cascade of events of metastasis of cells from an established tumour prior to their intravasation. To test definitively a potential role for YAP in tumour metastasis, we employed a spontaneous tumour metastasis model. MeWo cells expressing either a Dox-inducible control plasmid or one expressing YAP-5SA were engrafted into the flank of NOD/SCID Il2rγ-/- (NSG) mice, which lack an adaptive immune system, and tumours were allowed to grow until ~5mm in diameter (approximately 6 weeks). Half of the mice were then injected intraperitoneally with Dox for two consecutive days, and provided Dox in the drinking water until sacrificed, to induce expression of YAP-5SA. Mice were sacrificed when tumours reached 20mm in size, primary tumours harvested and metastasis to lymph nodes and secondary organs assessed (Fig. 4A).

**Figure 4.**
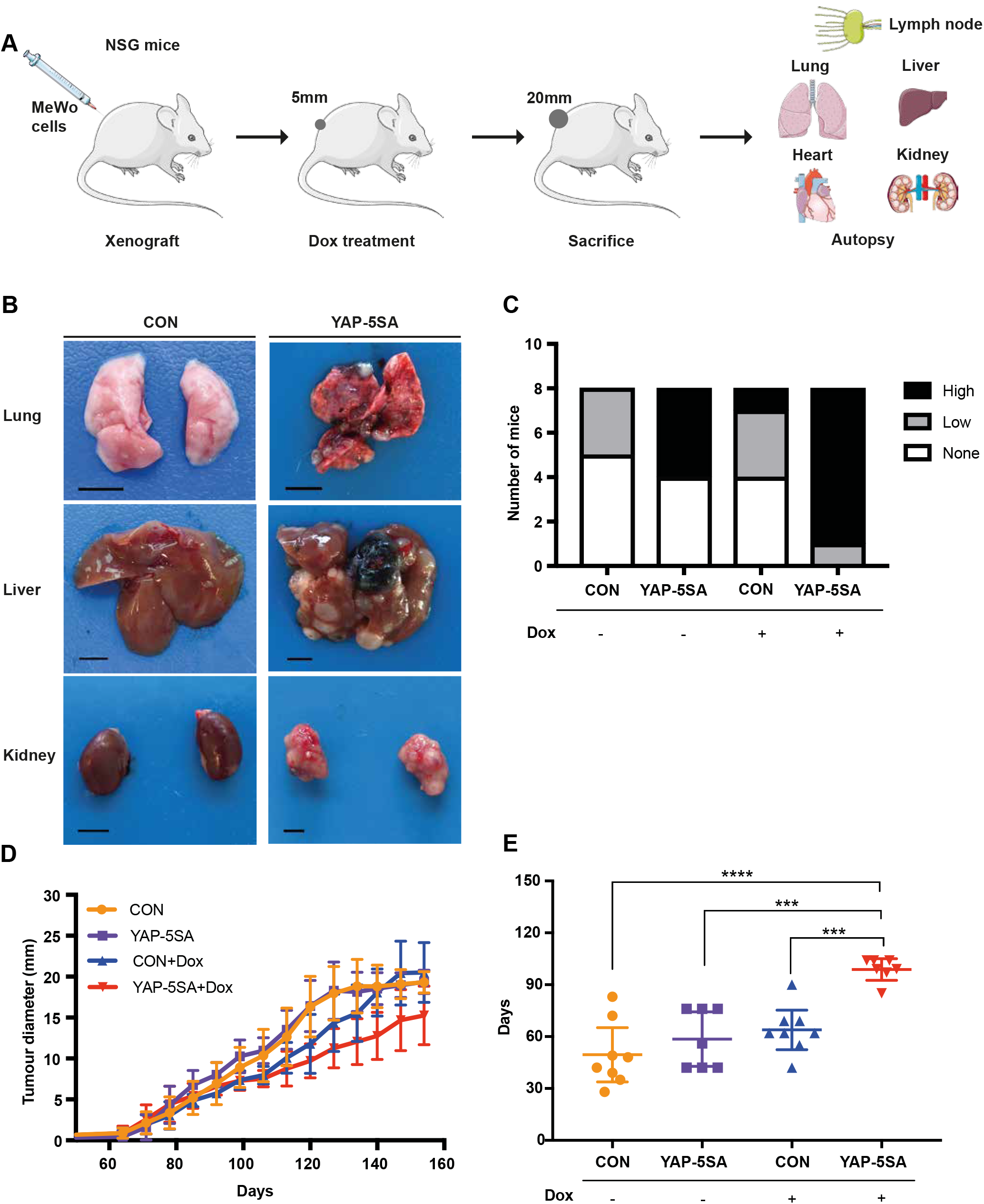
YAP induces spontaneous melanoma metastasis in vivo and hinders primary tumour growth. **A)** Schematic diagram of the initial spontaneous in vivo metastasis experiment. **B)** Representative images of the indicated organs from mice that harboured a xenograft that expressed either a control plasmid or YAP-5SA and were treated with Dox. **C)** Number of mice from the different experimental cohorts that displayed macrometastases. These harboured a primary tumour xenograft that expressed either a control plasmid or YAP-5SA, and were treated with or without Dox (n=6 in each group). **D)** Quantification of primary tumour growth in mice from the different experimental cohorts. **E)** Time taken (days) for primary xenografted tumours to grow from 5mm to 20mm in mice from the different experimental cohorts. Data are presented as mean ± standard deviation. Data are presented as mean ± standard deviation. ***P<0.001, ****P<0.0001.

All mice that were induced to express YAP-5SA in tumor cells generated metastases, and 7 out of 8 mice carried heavy burdens of macroscopically evident metastases in organs such as the lung, liver, kidney, heart, and lymph nodes (Fig. 4B and C). By contrast, more than half of the mice from the three control groups showed no signs of tumour metastasis, suggesting that YAP drives melanoma metastasis *in vivo* (Fig. 4B and C). However, the growth of YAP-5SA-expressing tumours was slower than control mice, such that these tumours took a longer time than controls to reach 20mm in size (Fig. 4D and E). This raised the possibility that the observed increase in metastasis was caused by the prolonged growth time and consequently increased ‘time in mouse’ of YAP-5SA+Dox tumours, rather than by an intrinsically enhanced metastatic capacity.

To test this, we repeated this experiment but allowed primary tumours to grow to 10mm in size before treating with Dox and when the first tumours in any cohort reached 20 mm in size, we harvested all mice and assessed metastasis (Fig. 5A). We also increased the numbers of mice in the Dox-treated groups to 9 each and reduced the non-treated groups to 3 each. Consistent with the initial experiment, YAP-5SA expressing tumours grew slower as they were significantly smaller than control tumours when mice were harvested (Fig. 5B). Importantly, we observed substantially more spontaneous metastases in mice harbouring YAP-5SA expressing primary tumours. All 9 mice in the YAP-5SA+Dox group had macrometastases in multiple organs (lung, liver, kidney and lymph nodes), whilst only 2 out of 9 mice in Ctrl+Dox group displayed macrometastases and these were only present in the lung (Fig. 5C). Additionally, only 3 of 6 mice in the other two control groups harboured macrometastasis (Fig. 5C).

**Figure 5.**
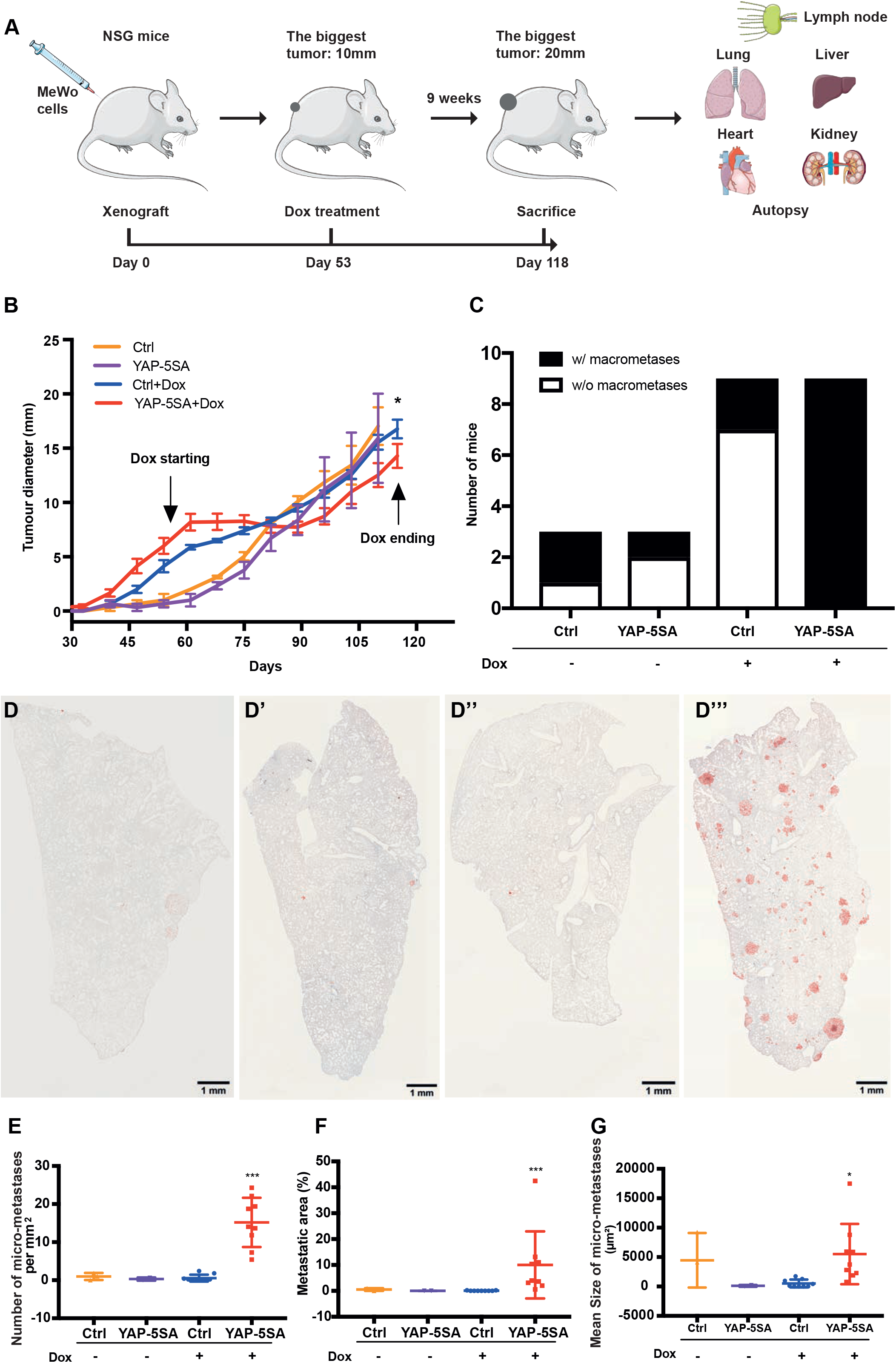
YAP induces spontaneous melanoma metastasis in vivo. **A)** Schematic diagram of the second spontaneous in vivo metastasis experiment. **B)** Quantification of primary tumour growth in mice from the different experimental cohorts. Data are presented as mean ± standard deviation. *P<0.05. **C)** Number mice that displayed macrometastases from the different experimental cohorts. These either harboured a primary tumour xenograft that expressed either a control plasmid or YAP-5SA and were treated with Dox (n=9 each), or did not receive Dox treatment (n=3 in each group). **D-D’’’)** Representative immunohistochemistry images of lung sections from mice that harboured a xenografted tumour that expressed either a control plasmid or YAP-5SA, and were treated with or without Dox. Lung sections were stained with anti-human mitochondria antibody (red stain) to reveal xenografted tumour cells that has metastasized. Scale bars = 1mm. **E-G)** Quantification of micrometastases in mice that harboured a primary tumour xenograft that expressed either a control plasmid or YAP-5SA and were not treated with Dox (n=3 each), and either a control plasmid or YAP-5SA and were treated with Dox (n=9 in each group). Quantifications were: the number of micrometasteses per mm2 (E), total metastatic area compared to non-metastatic area (F) and mean area of micrometasteses (G). Data are presented as mean ± standard deviation. *P<0.05, ***P<0.001.

As all mice in this second *in vivo* experiment were sacrificed when the biggest tumour reached 20 mm, the overall growth time of tumours was shorter than the first *in vivo* experiment, resulting in smaller metastases that were harder to distinguish with the naked eye (data not shown). To more accurately analyse metastasis in these mice, immunohistochemistry was performed to assess micrometastases. Given that lungs displayed the most macrometastases, these organs were sectioned and stained with an anti-human mitochondria antibody to identify xenografted melanoma cells that has metastasized to this organ (Fig. 5D-D’’’). In the YAP-5SA expression group, lung micrometastases were observed in all 9 mice, whilst only 2/3 of Ctrl/YAP-5SA-Dox mice, and 5/9 of Ctrl+Dox mice harboured lung micrometastases. Further, the number of micrometastases, the mean size of each micrometastatic lesion and the total micrometastatic area, were all substantially higher in YAP-5SA+Dox mice than in the three different control cohorts (Fig. 5D-G). These results indicate a strongly enhanced metastatic burden in YAP-5SA expressing mice, providing definitive evidence that YAP can promote spontaneous metastasis of melanoma cells *in vivo*.

### Identification of YAP target genes that mediate its ability to stimulate melanoma cell invasion

To identify potential target genes of YAP that mediate its ability to drive melanoma invasion and metastasis, we compared our melanoma YAP signature with the invasive melanoma signature previously reported by Hoek et al., (10) and found that ten genes were common to both signatures. We then determined whether these genes were present in one or more YAP signatures defined by the Piccolo laboratory in other cell lines and tissues (39–41) which allowed us to refine our list to six genes: *AXL, THBS1, CYR61, CRIM1, AMOTL2* and *FST*. *AMOTL2* is part of the Hippo pathway and is thought to repress YAP and be transcriptionally induced as part of a negative feedback loop (42). *FST* codes for a protein that binds to Activin and antagonizes its ability to contact TGF-β receptors (43); in the context of cancer it has been reported as an inhibitor of metastasis (44, 45). *AXL, THBS1, CYR61* and *CRIM1*, have all been reported to promote metastasis and/or cancer cell invasion and migration. *AXL* encodes a receptor tyrosine kinase and elevated *AXL* expression is linked to poor prognosis of melanoma, as well as contributing to invasion and metastasis in MITF-deficient melanomas (13, 14, 46). *AXL* is also a defining gene of the invasive melanoma cell state, and its expression is inversely correlated with MITF (13, 14, 46). *THBS1* encodes Thrombospondin 1, an adhesive glycoprotein that mediates cell-cell and cell-matrix interactions and melanoma cell invasion (47). *CYR61* encodes cysteine-rich angiogenic inducer 61 is a well-characterised YAP target gene, and has been implicated in different aspects of tumorigenesis including metastasis. *Cysteine-rich motor neuron 1 protein* (*CRIM1*) has been studied in the context of cancer cell migration and invasion, although conflicting results have been reported (48, 49). We thus tested potential roles for these four genes in cell invasion in vitro.

Initially, we tested the dependence of each of these genes on the invasive ability conferred to MeWo cells by YAP-5SA expression. Each gene was depleted by siRNA in YAP-5SA expressing MeWo cells and cell invasion compared to control siRNA. Depletion of AXL and THBS1 both strongly suppressed YAP-5SA-induced MeWo cell invasion (Fig. 6A, quantified in Fig. 6C). By contrast depletion of *CYR61* or *CRIM1* did not suppress YAP-induced invasion, although the invasion of *CYR61*-depleted cells trended towards significance (p=0.07) (Fig. 6A, quantified in Fig. 6C). We also investigated the role of the well-known YAP partner transcription factors TEAD1-4 in YAP-induced invasion, given that Verfaillie et al. used unbiased genomic approaches to link TEADs to the invasive melanoma cell state and showed that they were required for melanoma cell invasion in vitro (11). TEAD1-4 depletion almost completely blocked the ability of YAP-5SA-induced invasion of MeWo cells (Fig. 6A, quantified in Fig. 6C). siRNA-mediated depletion of each protein was confirmed by Western blot (Fig. 6B). Importantly, depletion of these different proteins did not impact cell viability within the time-course of these assays (Fig. 6D).

**Figure 6.**
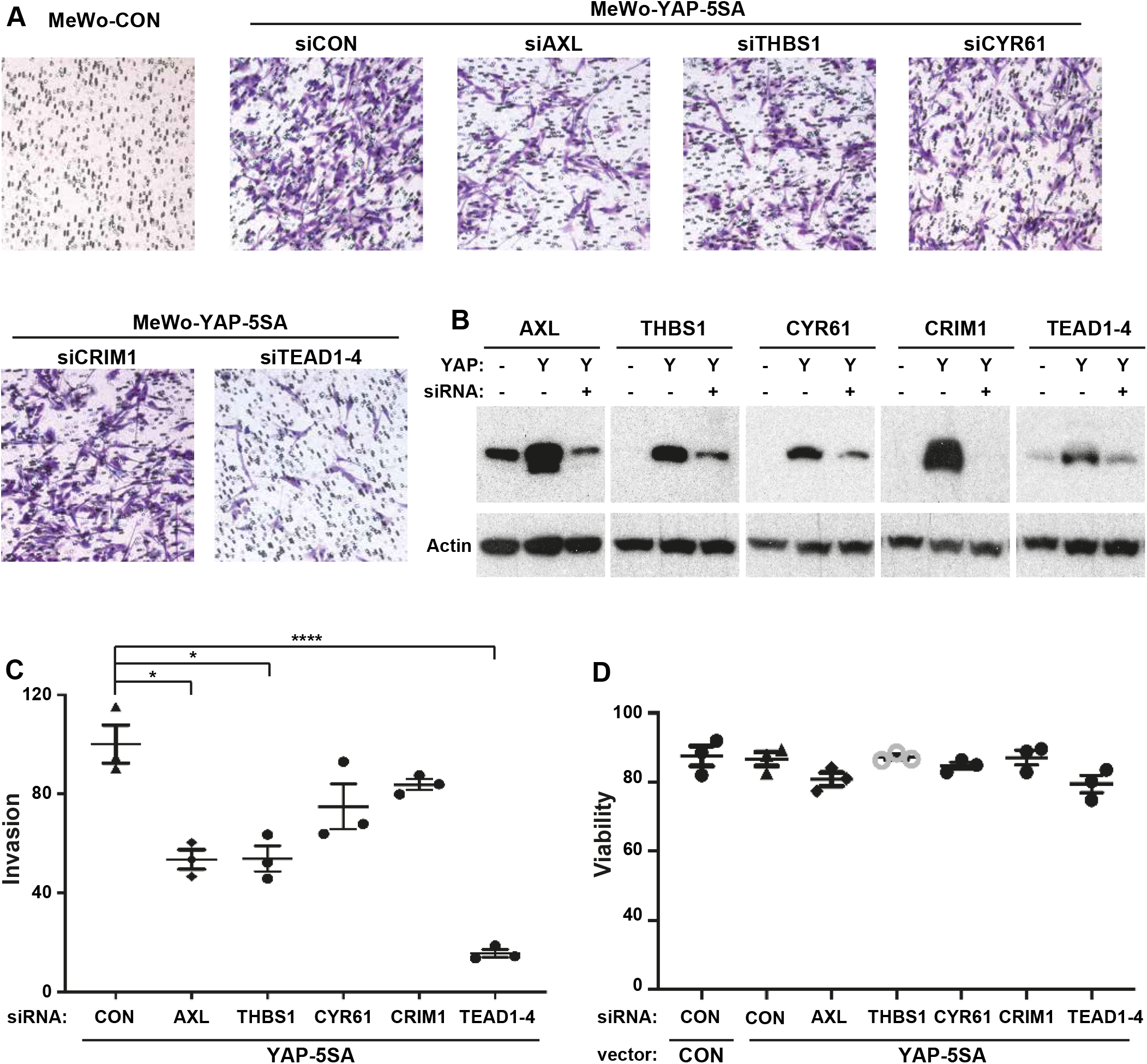
The YAP target genes *AXl*, *THBS1* and *CYR61* are required for melanoma cell invasion. **A)** Representative images of MeWo cells that were treated with Dox to induce YAP-5SA expression and treated with the indicated siRNAs and following invasion assays. **B)** Detection of AXL, THBS1, CYR61, CRIM1, TEAD1-4 and actin by western blot for the different siRNA treatments in (A). **C)** Quantification of invasion assays for the different siRNA treatments in (A). **D)** Quantification of cell viability assays for the different siRNA treatments in (A). Data in (C and D) are represented as mean +/- SEM from 3 biological replicates. *p<0.05, ****p<0.0001 (unpaired two-tailed t-tests or ANOVA plus Tukey’s multiple comparison tests).

Next, we assessed whether depletion of these YAP target genes affected the invasive ability of a melanoma cell line that has inherent invasive ability, elevated YAP activity and requires YAP for cell invasion. Both *AXL* and *THBS1* were required for C067-M1 cell invasion, as was *CYR61*. By contrast, *CRIM1* was not (Fig. 7A, quantified in Fig. 7C). Consistent with the studies of Verfaillie et al. (11), *TEAD1*-*4* depletion substantially hindered the invasive capacity of C067-M1 cells (Fig. 7A, quantified in Fig. 7C). siRNA-mediated depletion of each protein was confirmed by Western blot (Fig. 7B). As in our previous cell invasion experiments, depletion of these proteins did not affect cell viability in the time course of this experiment (Fig. 7D). Therefore, YAP mediates melanoma cell invasion by regulating expression of genes encoding for AXL and THBS1 and possibly also CYR61.

**Figure 7.**
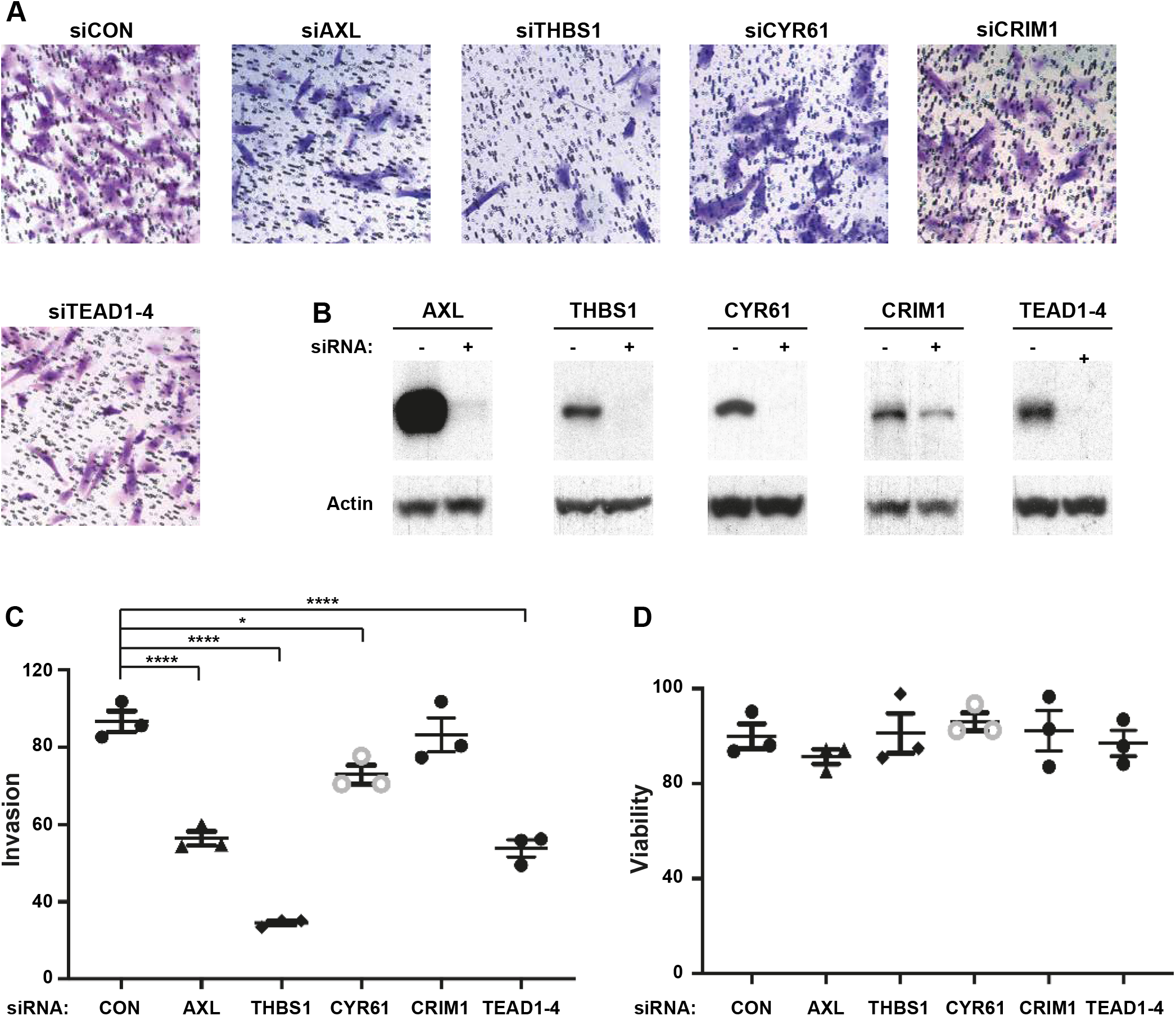
Identification of YAP target genes that mediate its ability to stimulate melanoma cell invasion. **A)** Representative images of C067-M1 cells treated with the indicated siRNAs and following invasion assays. **B)** Detection of AXL, THBS1, CYR61, CRIM1, TEAD1-4 and actin by western blot for the different siRNA treatments in (A). **C)** Quantification of invasion assays for the different siRNA treatments in (A). **D)** Quantification of cell viability assays for the different siRNA treatments in (A). Data in (C and D) are represented as mean +/- SEM from 3 biological replicates. *p<0.05, ****p<0.0001 (unpaired two-tailed t-tests or ANOVA plus Tukey’s multiple comparison tests).

## DISCUSSION

The Hippo pathway is an important tumour suppressor network in several human cancers although its role in cutaneous melanoma is less clear (23). Here, using a melanoma YAP transcriptional signature, we found that YAP activity is strongly enriched in invasive melanoma cell lines and is a driver of the invasive behaviour of these cells. Aligned with this, YAP drove melanoma metastasis in vivo in a spontaneous murine metastasis model. Previously, YAP was found to stimulate melanoma tumour seeding post tail vein injection (37, 38), but a role for YAP in metastasis had not been tested in a bona fide spontaneous metastasis model. Ours is thus the first demonstration that YAP hyperactivity promotes melanoma metastasis spontaneously from established tumours and raises the possibility that YAP could drive metastatic spread in patients. In accordance with this, Yap depletion limited metastasis of primary tumours to lymph nodes in a B16 murine melanoma model (50).

We found that YAP’s ability to stimulate melanoma cell invasion is dependent on TEAD1-4 transcription factors, which is in accordance with the studies of Verfaille et al., who utilised unbiased genomic approaches followed by cell based assays to identify a role for TEADs in melanoma cell invasion (11). We extend these studies by showing that YAP drives melanoma cell invasion by driving expression of genes encoding *AXL, THBS1* and *CRIM1*. From a therapeutic perspective, targeting of the physical interaction between YAP and TEADs is being keenly pursued as an anti-cancer strategy (51); compounds that disrupt this interaction would be predicted to limit both cancer growth and metastasis. Given that small molecule disruption of protein-protein interactions is challenging, another potential mode by which YAP’s pro-metastatic capacity could be blunted is by targeting one or more of its downstream target genes that mediate this activity, like AXL.

A conundrum in our studies was the finding that, while induced YAP hyperactivity promoted melanoma metastasis, it compromised the growth of primary tumours. Superficially, this finding runs counter to studies that identified a pro-growth role for the *Drosophila* YAP orthologue Yorkie (52), and subsequently, a similar role for YAP in many vertebrate organs, most notably the liver (fish and mice) (53–55). However, YAP hyperactivity does not always lead to tissue overgrowth. For example, in murine breast tissue, Yap hyperactivity caused by *Sav1* loss led to a terminal differentiation defect but not overgrowth (56). In murine cancer models, syngeneic xenografts that tested deletion of the *Lats1* and *Lats2* genes, which causes Yap hyperactivity, compromised tumour growth because an adaptive immune response was invoked (57). The latter finding cannot explain our results however because our experiments were performed in immunocompromised mice that lack an adaptive immune system. Instead, our data raise the interesting possibility that YAP-5SA expression drove metastasis but compromised primary tumour growth because forced constitutive YAP hyperactivity switches melanomas cells from a proliferative state to an invasive state while compromising their ability to switch between these states.

Single-cell sequencing and immunofluorescence studies have shown that melanomas consist of both invasive and proliferative cell types in differing ratios (12). Further, it is thought that melanoma cells are plastic and can oscillate between these two states by modulating their chromatin states and gene expression profiles. This plasticity is thought to facilitate their abilities to adapt to, grow and survive in different regions of the body and in changing tumour microenvironment. It has also been linked to response to therapy; for example, resistance to MAPK pathway inhibitors is causally linked to the invasive melanoma cell state (24–26). We propose that melanoma cells with constitutively high YAP in our experiments assumed the invasive state but lacked the ability to switch back to the proliferate state and so either died or downregulated YAP-5SA transgene expression to increase the chance of survival. This finding could provide an explanation for why mutations that promote constitutive YAP hyperactivity are rare in most cancers including melanoma (23). One prediction from this is that YAP hyperactivity would peak in cells prior to metastasis to drive this phenomenon but then be downregulated to favour survival and tumour development from metastasized cells.

Our finding of the key role of Hippo signalling in regulating melanoma cell invasion in vitro and spontaneous metastasis in vivo highlight the importance of identification and development of inhibitors of YAP activity. Such therapeutics would be predicted to protect cancer patients from complications of the metastatic process, which contribute the vast bulk of morbidity and mortality conferred by malignant disease.

## MATERIALS AND METHODS

### Cell Culture

Melanoma cell lines were cultured in RPMI 1640 + 20mM HEPES medium (Gibco), supplemented with 10% fetal bovine serum (FBS) (Gibco) and 1% Penicillin Streptomycin (P/S) (Gibco). For siRNA transfections, cells were seeded into 6-well plates 24 hours (hrs) before transfection. Media were removed and replaced with P/S-free fresh RPMI media, Lipofectamine RNAiMAX transfection reagent (Invitrogen), 5μM siRNAs and 100 μl of Opti-MEM Reduced Serum Media (Gibco) for 5 minutes (min) at room temperature (RT). For overexpression studies, cell lines that stably expressed Dox-inducible plasmids were generated by retrovirus transduction as in (36), and treated with between 0.03-0.1µg/ml Dox. Cells were trypsinised 24 hrs after siRNA or Dox treatment for use in various assays or collected 48 hrs later for immunoblots.

### Cell Invasion and Migration Assays

Millicell Hanging Cell Culture Insert with PET 8 µm transwell inserts (Merck) were placed into 24-well plates and coated overnight with 10μg of Matrigel (Corning) diluted in 100μl of serum-free RPMI media at 37°C. 6×10^4^ melanoma cells per insert were added to the insert of each chamber in 200μl of serum-free RPMI media. The lower chamber was loaded with 600μl of RPMI media supplemented with 10% FBS. After 48hrs, inserts were washed with PBS twice before and after staining with a 0.1% Crystal Violet solution for 15 min. Non-invaded cells remaining on upper layer of inserts were removed by cotton swabs. Invaded cells were imaged using an inverted microscope (Zeiss Axio Vert. A1; Carl Zeiss AG, Oberkochen, Germany) at a magnification of 100×. Five fields were randomly chosen for each insert and cells counted manually. Migration assays were performed as above but plates were not coated in matrigel.

### Cell Viability Assays

Cells were seeded in 96-well plates and transiently transfected with siGENOME SMARTpool siRNAs or OTP control siRNAs (ON-TARGETplus Non-Targeting Pool siRNA) using Lipofectamine® RNAiMAX transfection reagent (Invitrogen). Media was changed after 24hrs and cells incubated for a further 72 hrs. Alamar Blue was added to each well and incubated at 37°C for 2 hrs. Fluorescence was read using a POLARstar OPTIMA (BMG Labtech) at 540/610 nm.

### Immunoblotting

Whole cell lysates were prepared in RIPA buffer with protease and phosphatase inhibitor cocktails (Roche). Lysates were subjected to SDS PAGE electrophoresis and transferred onto PDVF membrane (Millipore). Membranes were probed with antibodies against YAP (Cell Signaling Technology 4912), Actin (Cell Signaling Technology 4967), AXL (Cell Signaling Technology 8661), THBS1 (Novus Biologicals NB-100-2059), CYR61 (Cell Signaling Technology 14479), CRIM1 (Sigma SAB3500847) or pan-TEAD1 (Cell Signaling Technology 13295) followed by horseradish-peroxidase conjugated secondary antibodies and chemiluminescent detection.

### RNA sequencing

Total RNA was collected from cultured melanoma cells using Trizol, according to the manafacturer’s protocol (Invitrogen) and RNA quantity checked using Qubit RNA HS (Thermo Fisher Scientific). 1mg total RNA was used for library preparation according to standard protocols (QuantSeq 3’ mRNA-Seq FWD, Lexogen). Indexed libraries were pooled and sequenced on a NextSeq500 (Illumina). 5-15 million single-end 75bp reads were generated per sample.

### Quantitative real-time polymerase chain reaction (qPCR) analysis

Reverse transcription was performed on total RNA using a SuperScript III Reverse Transcriptase kit (Invitrogen). QPCR was performed with the Fast SYBR^®^ Green Master Mix, according to the manafacturer’s protocol (Applied Biosystems). PCRs were run on a StepOne Plus instrument, using the StepOne Plus software (Applied Biosystems). Data were normalized to *GAPDH* expression.

### Bioinformatics

RNA sequencing data were processed using Seqliner RNA-Seq alignment pipeline (v0.4; seqliner.org). Reads were aligned to GRCh37/hg19 using Tophat 2 (58). Mapped reads were counted using HTSeq package (59) to obtain the read counts for each gene. The R package *‘limma’*(60) was used to perform differential expression analysis. Specifically, we performed *voom* normalization and then linear modeling of data to obtain differentially expressed genes between different conditions.

### Analysis of Melanoma Cell Line data

We analyzed gene expression data from 55 human metastatic melanoma cell lines (61), available from https://www.ebi.ac.uk/arrayexpress/experiments/E-MTAB-1496/. The mutation status for commonly mutated genes in melanoma (BRAF and NRAS) were retrieved from (61). Among *BRAF* mutations, there were 23 V600E, seven V600K, one K601E and two G469E. *NRAS* mutations were found in 11 cell lines; four were Q61K, two Q61H and five Q61Q. The transcriptomes from each cell lines were profiled using Illumina HumanHT-12 V4.0 expression beadchip. Raw data were pre-processed using R statistical software (https://www.r-project.org/) ‘*lumi’* package. Data were log2 transformed and quantile normalized. Hierarchical clustering was performed to identify groups of cell lines that displayed similar gene expression profiles to the YAP melanoma signature. First, we extracted normalized gene expression levels of each gene in the gene signature and calculated Pearson correlation between the expression levels. The pairwise dissimilarities were calculated as *1-correlation/2*. Then, the ‘*hclust’* function (R package stats) was used to perform hierarchical clustering (average method). Finally, heat maps were produced using the heatmap.2 function (R package gplots). Gene set enrichment analysis (GSEA) (62) was performed on the YAP melanoma signature and to obtain the ranked gene lists based on GSEA normalized enrichment scores.

### Animal Experiments

All experiments were carried out in accordance with the Peter MacCallum Cancer Centre Animal Ethics and Experimentation Committee protocols (#E526). NOD/SCID Il2rg-/- (NSG) mice were subcutaneously transplanted with GFP^+^ human melanoma MeWo-CON or MeWo-YAP-5SA cells to generate melanoma xenografts. Mice were randomly separated into control group and doxycycline (Dox) treatment group. The Dox-treated group received intra-peritoneal injection of 40ug of Dox per mouse for two consecutive days in addition to drinking water supplemented with dox at a concentration of 2 mg/ml until sacrificed. Tumour growth was monitored every week by palpation, and tumour diameter was measured with a vernier caliper. Growth rates were determined by maximum tumour diameter (in mm) divided by time elapsed (in weeks) from the date tumours first became palpable.

### Cell Preparation, Labelling and Flow Cytometry

Fluorescent Activated Cell Sorting was used to identify transduced cells that expressed GFP. Cultured cells were trypsinised and coated in PBS supplemented with 2% of FBS. For tumour cell preparation, tumours were mechanically dissociated with a McIlwain tissue chopper (Mickle Laboratory Engineering). Enzymatic tumour dissociations were performed according to published methods (63). Antibody labelling was performed for 30 minutes on ice. Cells were stained with directly conjugated antibodies to human HLA-A, B, C (1:5, G46-2.6-PE, BD Pharmingen), mouse CD45 (1:200, 30-F11-APC, BD Pharmingen), mouse Ter119 (1:100, TER-119-APC, BD Pharmingen), and mouse CD31 (1:100, 390-APC, eBioscience) to enable selection of HLA^+^CD45^−^ TER119^−^CD31^−^ (Lin-) GFP^+^ cells. Labelled cells were resuspended in 10ug/ml DAPI (Roche) for viability and analysed on a BD LSR II flow cytometer (BD Biosciences), or a BD FACS Canto II flow cytometer (BD Biosciences). Cell sorting was performed by technicians from the FACS Facility of Peter MacCallum Cancer Centre using the BD FACS AriaII Cell Sorter (BD Biosciences).

### Immunohistochemistry

Tumour and organ tissues were fixed in 10% neutral-buffered formalin (Australian Biostain) overnight at room temperature. The samples were embedded in paraffin by the Centre for Advanced Histology and Microscopy of Peter MacCallum Cancer Centre Paraffin sections of 4µM thickness were heated in an oven at 60°C for 30 min, dewaxed in histolene, and hydrated through graded alcohols and distilled water. Sections were subjected to heat-induced antigen retrieval in target retrieval solution (Dako) at 125°C for 3 min heated by a pressure cooker (Biocare Medical). The sections were allowed to cool down, and washed in TBS for 3 times. Quenching of endogenous peroxidase was performed in freshly made 3% hydrogen peroxide in methanol for 15 min. The sections were then sequentially washed in TBS for 3 times, incubated in blocking solution (1% BSA in TBST) for 60 min at RT. Sections were then incubated with mouse anti-human mitochondria (clone 113-1, Merck) diluted in blocking solution at a dilution of 1:1000 at 4°C overnight. After washing with TBST, the slides were incubated with secondary antibody using an ImmPRESS™ HRP Anti-Mouse IgG (Peroxidase) Polymer Detection Kit (Vector Laboratories) for 60 min at RT. Sections were washed with TBST and developed using AEC substrate-chromogen (Dako) for 5 min. The samples were counterstained with haematoxylin for 10 sec, washed with distilled water, and differentiated in Scott’s tap water for 30 sec. Sections were cover slipped with Aquatek (Merck) prior to imaging on an Olympus VS120 Virtual Slide Microscope. Metastasis analysis was performed based on positive staining of the human mitochondria on 10 random 20x fields per sample using FIJI (v1.49).

### Statistics

Statistical analysis was performed using GraphPad Prism version 7. Differences between mean tumour growth rates were compared using ANOVA followed by Tukey’s multiple comparison test. Other analyses were performed using unpaired Student’s t-test or ANOVA as appropriate. Tukey’s multiple comparison tests were used to compare individual groups.

## ACKNOWLEDGEMENTS

We thank the following Peter MacCallum Cancer Centre core facilities: Molecular Genomics, Bioinformatics, Flow Cytometry and the Centre for Advanced Histology and Microscopy, which were in part supported by the Australian Cancer Research Foundation. K.F.H was supported by a National Health and Medical Research Council (NHMRC) Senior Research Fellowship (APP1078220). M.S. was supported by Pfizer Australia, NHMRC, veski, and VCA Fellowships. This research was supported by grants from the Cancer Council of Victoria (APP1080255) and NHMRC (APP1145166), and by the Peter MacCallum Cancer Foundation.

